# Inferring temporal organization of postembryonic development from high-content behavioral tracking

**DOI:** 10.1101/2020.11.11.378166

**Authors:** Denis F. Faerberg, Victor Gurarie, Ilya Ruvinsky

**Author notes:** Correspondence to: Ilya Ruvinsky.

## Abstract

Understanding temporal regulation of development remains an important challenge. Whereas average, species-typical timing of many developmental processes has been established, less is known about inter-individual variability and correlations in timing of specific events. We addressed these questions in the context of postembryonic development in *Caenorhabditis elegans*. Based on patterns of locomotor activity of freely moving animals, we inferred durations of four larval stages (L1-L4) in over 100 individuals. Analysis of these data supports several notable conclusions. Individuals have consistently faster or slower rates of development because durations of L1 through L3 stages are positively correlated. The last larval stage, the L4, is less variable than earlier stages and its duration is largely independent of the rate of early larval development, implying existence of two distinct larval epochs. We argue that characteristic patterns of variation and correlation arise because duration of each stage tends to scale relative to total developmental time. This scaling relationship suggests that each larval stage is not limited by an absolute duration, but is instead terminated when a subset of events that must occur prior to adulthood have been completed. The approach described here offers a scalable platform that will facilitate the study of temporal regulation of postembryonic development.

## Introduction

As is true for other Ecdysozoa (Telford et al., 2008), postembryonic development of nematodes is organized into several discrete stages, separated by molts. Upon completing embryonic development, *C. elegans* progress through four larval stages (L1-L4) prior to larval-to-adult transition (Byerly et al., 1976). Between L1 and adulthood, freely moving larvae execute stage-specific developmental programs that increase the total (in hermaphrodites) number of somatic nuclei from 558 to 959 (Sulston et al., 1983), produce ~2,500 germline nuclei (Hirsh et al., 1976), while allowing the worms to grow on average from ~250 to ~1,000 μm in length (Byerly et al., 1976; Hirsh et al., 1976).

Larval stages have similar organization – the multi-hour periods of growth are capped by short periods of ecdysis, during which the old cuticle is shed (Singh and Sulston, 1978). Particular developmental events (e.g. cell divisions, deaths, migration, etc.) occur at specific times (Sulston and Horvitz, 1977) and transitions between larval stages are characterized by dramatic upheavals in gene expression (Frand et al., 2005; Hendriks et al., 2014; Snoek et al., 2014; Turek and Bringmann, 2014). Genetic analysis of timing of developmental events led to discovery of heterochronic mutants (Ambros and Horvitz, 1984), including the now-classic miRNAs *lin-4* (Lee et al., 1993) and *let-7* (Reinhart et al., 2000), as well as their targets (Slack et al., 2000; Wightman et al., 1993), and other genes (Abbott et al., 2005; Abrahante et al., 2003; Antebi et al., 1998; Jeon et al., 1999; Monsalve et al., 2011; Moss et al., 1997; Rougvie and Ambros, 1995) that regulate timing of developmental transitions (Rougvie and Moss, 2013).

Approximate population-average timeline of development is sufficient for analysis of the overall order of events; these estimates were made in the early days of *C. elegans* research (Byerly et al., 1976; Hirsh et al., 1976), but remain relevant today. They do not, however, permit inferences of inter-individual variation of developmental rates or more involved analyses of co-dependence of different developmental events and stages. Direct observation of developmental progression is time-demanding and labor-intensive, necessarily limiting numbers of animals that can be followed simultaneously. High-throughput approaches relying on a variety of technologies have been developed (Gritti et al., 2016; Keil et al., 2017; Nika et al., 2016; Olmedo et al., 2015; Uppaluri and Brangwynne, 2015), including methods that allow long-term observation (Stroustrup et al., 2016; Zhang et al., 2016). Some of these platforms could in principle be used to analyze progression of development in individual animals. One promising approach is based on the periodic nature of the locomotor activity during postembryonic development – episodes of ecdysis at the end of each larval stage are preceded by periods of lower activity, called lethargus, that last ~1-2 hours (Singh and Sulston, 1978). Therefore, identifying periods of lower activity from continuous recordings could yield estimates of larval stage duration (Raizen et al., 2008), even though individuals are not uniformly inactive during lethargus (Iwanir et al., 2013). Recently, Stern *et al.* reported behavioral analysis of several hundred continuously monitored singled hermaphrodites over a period that extended from the onset of L1 to beyond the L4/adult transition (Stern et al., 2017). Taking advantage of these data, we set out to assess inter-individual variability and relationships between different stages of postembryonic development.

## Results

### High-content behavioral tracking data can reveal temporal progression of development in individual *C. elegans*

To investigate long-term behavioral patterns in *C. elegans* hermaphrodites, Stern *et al.* continuously monitored individuals singled from hatching and freely moving on hard agar surfaces within relatively large (~10 mm, *i.e.* >10 times larger than the length of larvae) arenas (Stern et al., 2017). An advantage of relying on these data to ascertain larval stage duration, compared to more invasive methods or ones that restrain larvae during development, is that in this paradigm larvae moved freely and experienced minimal disturbance. Animals were observed from the onset of movement during early L1 stage until early adulthood, in the presence of *E. coli* food. Each of the 125 individuals in the study was imaged for >50 hours at a frequency of 3 frames per second resulting in ~6 × 10^5^ frames/worm.

Although complete tracks generated over the entire duration of a recording were highly convoluted, coordinates of “centers of mass” captured in adjacent frames (*i.e.*, 1/3 seconds apart) could be used to compute a quantity that characterizes animals’ movement; we refer to this quantity as displacement (Figure 1A). Plotting all (~6 × 10^5^) sequential displacements, provides a dynamic picture of movement activity over the entire duration of larval development (Figure 1B). Such plotting alone could reveal approximate periods of lower activity, at least in some individuals. Although some periods of low activity likely reflect lethargus episodes surrounding molts (Raizen et al., 2008), displacements vary considerably between frames (Figure S1A), making it challenging to computationally identify periods of lower activity, particularly in some individuals (Figure S1B). To overcome these limitations and to leverage the power of inter-individual comparisons, we implemented a method for identifying periods of lower activity from the somewhat noisy activity data (like those shown in Figure 1B). Typical velocity in the presence of food is ~30-100 μm/s (Ramot et al., 2008) for larvae that range between ~250 and ~1,000 μm (Byerly et al., 1976; Hirsh et al., 1976). We therefore reasoned that displacements over 1/3 sec (in the data we analyzed, these averaged ~4-5 μm) largely reflect minor changes in body posture, including head movements (Nagy et al., 2014; Yemini et al., 2013), rather than genuine locomotion. Following extensive testing, we determined that selecting 1 out of every 30 frames represented a reasonable compromise between de-noising the data and sampling sufficiently frequently to capture movement patterns. This level of data reduction is equivalent to recording activity at 1 frame per 10 sec, which has been empirically found to be an appropriate frequency based on different considerations (Huang et al., 2017; Nelson et al., 2013; Raizen et al., 2008). Because the 30X-reduced activity profiles were still quite noisy (Figure S1C), we tested whether sliding windows of various length could generate smoother curves without removing features essential for identifying periods of lower activity. We found windows of ~333 frames of 30X-reduced activity (10,000 frames of primary, unreduced data) to be useful for this task; such frames cover ~55.5 minutes of developmental time. Using these data reduction and smoothing parameters, we generated activity profiles for all 125 individuals in the data set. In every case, the profiles had four well-articulated periods of lower activity that by timing and duration approximately corresponded to lethargus periods (see below). In all 125 activity profiles we identified mid-points within periods of lower activity and designated corresponding times as boundaries between adjacent larval stages (Figure 1C).

**Figure 1.**
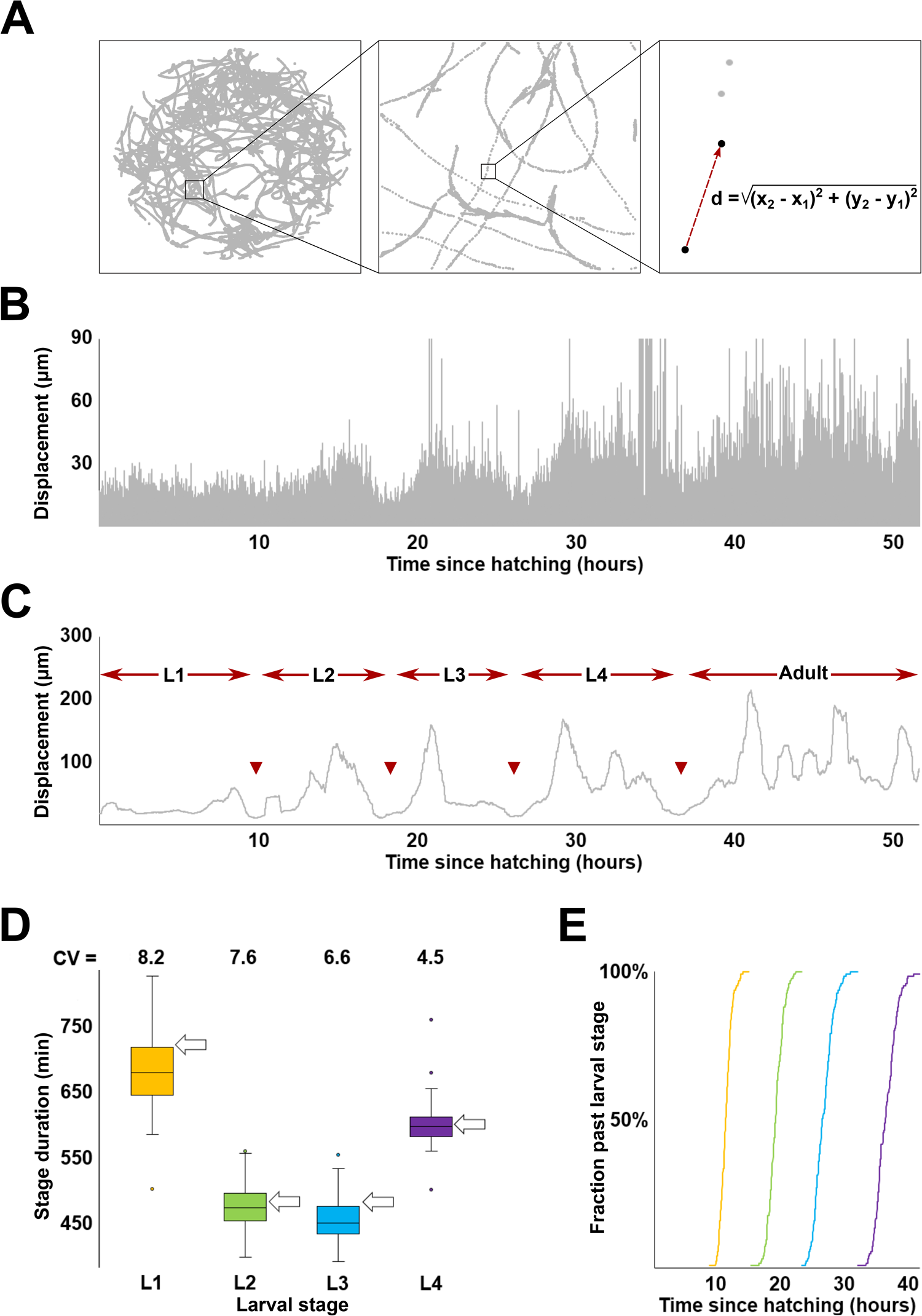
Inferring duration of larval stages from high-content behavioral tracking data. **(A)** Track of a single *C. elegans* hermaphrodite over the course of an ~50 hour recording. The right-most box shows calculation of displacement as distance between centers of mass of the tracked worm between two sequential frames. **(B)** Activity profile (*i.e.*, plot of all consecutive displacements) of the worm shown in (A). Note that due to fluctuations in locomotor behavior, the ~600,000 displacements shown here exaggerate local extremes; vast majority of displacement values are considerably lower (mean ~4-5 μm) than the outline. See Figure S1 for more detail. **(C)** Activity profile of the worm shown in (A) produced from displacement values sampled at 0.1 Hz and smoothened (55.5 min). Arrowheads indicate boundaries between larval stages defined as mid-points of periods of reduced activity. **(D)** Inferred durations and coefficients of variation (CV; expressed as %) of L1-L4 larval stages. Arrows indicate stage durations (at 22°C) shown in (https://www.wormatlas.org/hermaphrodite/introduction/Introframeset.html). **(E)** Transition times (sample N=125) between (L1, L2, L3, L4) larval stages.

Having corrected for an artifact of window smoothing and missing frames (see Materials and Methods), we obtained estimates of durations of larval stages (Figure 1D). These estimates closely matched those previously obtained from direct observations (Gritti et al., 2016; Hirsh et al., 1976; Monsalve et al., 2011). Discrepancies were minor (<1 hour compared to 8-11 hour stage durations) and could be due to rounding of prior estimates, to minute differences in cultivation conditions (e.g., between 20°C and 25°C, temperature increase of 1°C accelerates larval development by ~2 hours (Gouvea et al., 2015)), or to other difficult-to-control factors. Our estimates matched well those previously made from the same data (Stern et al., 2017), while the fractions of overall developmental time occupied by L1-L4 were virtually identical between our analysis and that of Raizen *et al.* (Raizen et al., 2008), even though recordings were conducted at different temperatures (see more on this below). Durations of individual larval stages could be used to compute transition times between larval stages for the entire population (Figure 1E). Despite the relatively modest variation overall (coefficient of variation (CV) of timing of L4/adult transition is ~4.6%), the slowest developing individual reached adulthood ~10.1 hours later than the fastest, a considerable difference given the ~36.9 hour average duration of larval development.

### Developmental rate is largely decoupled from behavioral activity

We tested whether our estimates of duration of larval stages were correlated with the locomotor activity data from which they were derived. We found at best a modest correlation between duration of larval development and activity, which was computed as sum of displacements divided by time (Figure S2A; see Materials and Methods). Duration of the L2 stage was correlated with activity, while L4 showed marginal correlation and L1 and L3 stages showed none (Figure S2B). It is possible that locomotor activity *per se* is not the appropriate measure to evaluate correlation between behavior and duration of larval development. We therefore tested whether fraction of time devoted to roaming, a related but distinct measure (see Materials and Methods) was better suited for the task. We found that the overall correlation was slightly higher, with only L2 (and possibly L4) showing evidence of correlation (Figure 2A, B).

**Figure 2.**
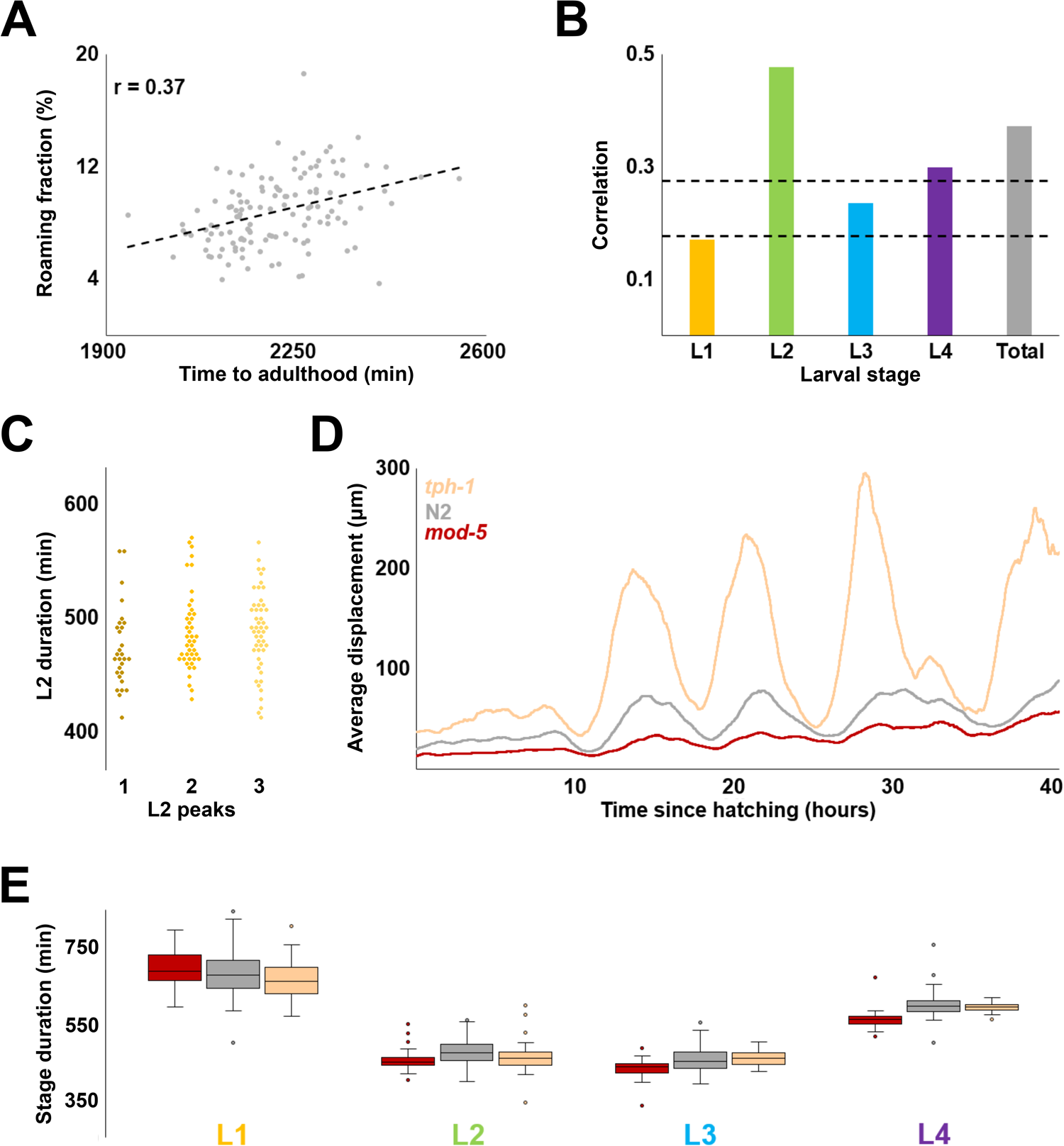
Relationship between the rate of development and locomotor activity. **(A)** Correlation between total developmental time and locomotor activity (measured as roaming fraction) for 125 wild type N2 worms. **(B)** Correlation between roaming fraction and stage duration for each larval stage; “total” shows the same value as in (A). Dashed lines denote two and three standard deviations above the expected correlation between two random sets of 125 uncorrelated variables. **(C)** Durations of L2 stages from each of the three categories activity profiles classified by overall shape (each diamond is one individual). Of the three possible pairwise comparisons (Kolmogorov-Smirnov test) only one – 1 peak vs 3 peak – had a p-value < 0.017 (0.05 after the Bonferroni correction for 3 comparisons is 0.017). The observed value – 0.015 – indicated at best a marginal difference. **(D)** Population average activity profiles of *tph-1(mg280)* (N=47), wild type N2 (N=125), and *mod-5(n822)* (N=41). **(E)** Inferred stage durations of *tph-1,* N2, and *mod-5* animals.

As we examined activity profiles of individual animals, we noticed that despite diversity of shapes, these profiles displayed repeated patterns of higher and lower activity within larval stages, often in stereotyped, albeit complex ways (Figure S2C). We reasoned that shapes of activity profiles may reflect some currently unknown feature(s) of behavior. If so, it is possible that individuals displaying different activity profiles might develop at different rates. To test this idea, we focused on the L2 stage because we expected it to offer the best chance of identifying a relationship, if one exists, between activity and duration of development. We manually classified the 125 animals in the data set into one of three categories by the shape of L2 activity profiles. We found no appreciable differences between the three categories with respect to the duration of L2 stages (Figure 2C).

The method described here could be used to analyze temporal unfolding of larval developmental in mutants (Figure S2D). Even in cases of noisy activity profiles (Figure S2E), we were able to infer total duration of larval development (Figure S2F). Our estimates are consistent with the ones made previously (Stern et al., 2017). As an additional test of whether there exists a relationship between locomotor activity and duration of larval development, we examined two mutants (Figure 2D). The first is in the *tph-1* gene that encodes a serotonin biosynthetic enzyme tryptophan hydroxylase and is consequently defective in serotonergic signaling. The second mutation affects the serotonin reuptake transporter gene *mod-5*, effectively increasing serotonergic signaling. Compared to wild type, these two mutant strains are known to have exacerbated and reduced exploratory behavior, respectively (Flavell et al., 2013; Stern et al., 2017). Consistent with prior analysis (Stern et al., 2017), despite having an approximately five-fold difference in activity (Figure S2G), the two mutants have nearly indistinguishable average durations of all larval stages (Figure 2E). It remains formally possible that *tph-1* and *mod-5* mutations proportionally scale both developmental rate and activity (see more on this below). Still, the most plausible interpretation of these results is that no simple relationship exists between activity and rate of larval development.

### Inferring temporal organization of larval development

The principle advantage of well-resolved developmental time series obtained from individual animals, compared to population averages, is that they could be used to explore inter-individual variability and relationships between different larval stages. Durations of L1-L4 larval stages were distributed approximately normally, with only a small number (~3 out of 125) of extreme outliers (Figure 3A). Same was true for total (*i.e.*, L1+L2+L3+L4) durations of larval development (Figure 3B). We wondered whether animals that developed much slower or much faster than average did so because of one or two abnormally fast or slow larval stages. We therefore compared durations of all larval stages between 20 animals with the fastest and 20 animals with the slowest development. We found that the two populations had nearly nonoverlapping distributions of L1 through L3, but indistinguishable L4s (Figure 3C). Largely the same relationships were observed when the fastest and slowest individuals were excluded from the analysis (Figure S3A). We interpret these results as an indication that A) animals develop at characteristic rates that are somewhat stable during the first three larval stages and B) the duration of the L4 is independent of those rates.

**Figure 3.**
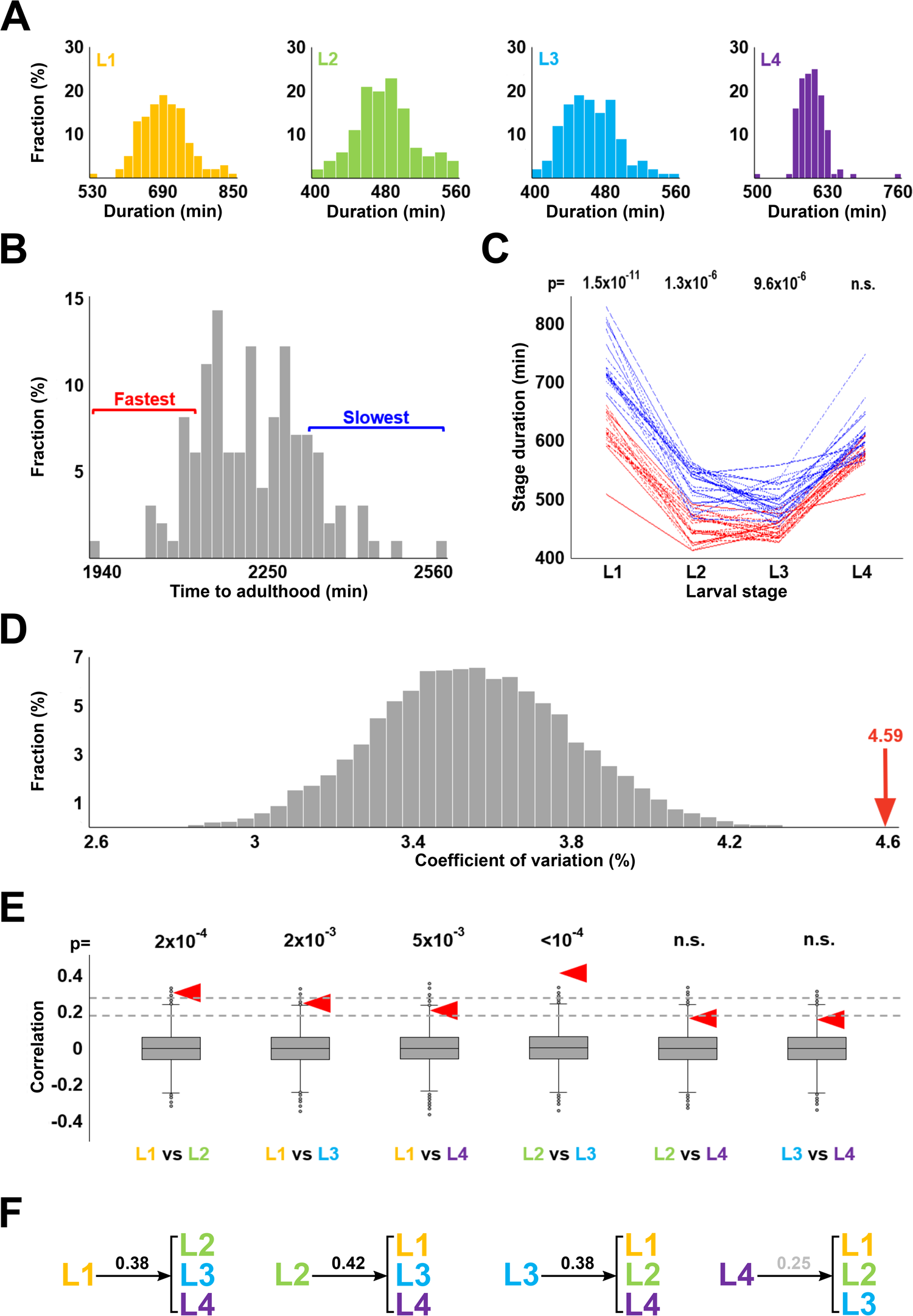
Correlations between stage durations. **(A)** Histograms of stage durations of wild type N2 animals (N=125). Durations of L1, L2, and L3 stages (as well as total durations of development; not shown) are consistent with being sampled from normal distributions (according to Shapiro-Wilk tests). Durations of L4 could be made normal if as few as 3 extreme outliers were removed. **(B)** Histogram of total developmental times of wild type N2 animals (N=125). Red and blue brackets denote 20 fastest and slowest developmental times, respectively. **(C)** Stage durations of the 20 fastest and 20 slowest developing worms (by time to reach adulthood). p-values shown above each stage are results of the Kolmogorov-Smirnov test comparing durations of that stage for the 20 fastest and 20 slowest developing worms. **(D)** Coefficients of variation of total duration of development in 10,000 sets of 125 artificial developmental time series constructed from randomly selected stage durations. Red arrow represents the coefficient of variation of the total durations of development of the 125 empirical activity profiles. **(E)** Correlation of stage durations in 10,000 artificial data sets. Null hypothesis is that compared variables are independent, and thus their correlation is 0. Dashed lines denote two and three standard deviations from this expected correlation. Red arrowheads show correlation values obtained from the empirical dataset. p-values above each comparison are the fractions of instances (out of 10,000) in which CVs of randomly permuted data are greater than those from the empirical dataset. **(F)** Correlation coefficients between duration of an indicated larval stage and the duration of the remaining three stages. Correlation coefficient for L4 is shown in grey because it is lower than three standard deviations from the expected correlation of two random sets of variables (N=125).

To systematically explore the apparently nonrandom associations between larval stage durations, we computationally generated 10,000 data sets, each containing 125 combinations of L1, L2, L3, and L4 stage durations that were randomly selected from respective empirical data. Each combination of L1-L4 simulated a developmental time series of an individual animal, while each set of 125 combinations matched in size the empirical data set analyzed here.

For each set of the 125 simulated developmental time series we computed coefficient of variation (CV) of total larval development times. We thus obtained 10,000 values of CVs on the assumption that each developmental time series is randomly assembled from an L1, an L2, an L3, and an L4 (Figure 3D). Because the CV of the empirical data (4.59) is considerably greater than expected on the assumption of random association (p<10^−4^), correlations must exist between stage durations. Of the six possible pairwise comparison of the four larval stages, two – L1 vs L2 and L2 vs L3 – showed significant positive correlation (Figure 3E), although only the latter remained significant when extreme values were removed from consideration (Figure S3B). In addition, we observed correlation between duration of the L1 stage and the remainder of larval development (L2+L3+L4); same was true for L2 and L3, but not the L4 (Figure 3F). In all comparisons, correlations involving the L4 tended to be lower than those involving other stages.

### Fractional scaling of developmental time series

One possible mechanism that could control duration of postembryonic development in *C. elegans* is “absolute timer” that allots specific time to each larval stage, variation being a consequence of intrinsic and extrinsic noise. Although normal distributions of L1-L4 (Figure 3A) and total (Figure 3B) durations would be expected under this scenario, correlations we detected between stages (Figure 3C-F) are inconsistent with the strict version of the model. Instructively, our estimates of the fractions of total time of postembryonic development devoted to L1-L4 (0.31, 0.21, 0.21, 0.27, respectively) are indistinguishable from those obtained in several independent studies that relied on different methodologies and were conducted at different temperatures (Byerly et al., 1976; Gritti et al., 2016; Hirsh et al., 1976; Keil et al., 2017; Raizen et al., 2008). We therefore studied “fractional times” that can be computed from absolute times – for each individual, fractional duration of a stage is equal to absolute duration of that stage divided by total developmental time.

In the empirical set of 125 individuals, fractional times were considerably less variable than the absolute times from which they were derived (compare CVs in Figure 4A vs Figure 1D). We next tested whether the CVs of fractional stage durations obtained from the empirical data set were lower than would be expected if stage durations in each individual were randomly associated. We examined variability of fractional stage durations of the 10,000 randomized sets (same sets as analyzed in Figure 3) and found that empirical data were less variable, particularly in the L2 and L3 stages (Figure 4B). Is variability of absolute stage durations always higher than the variability of fractional stage durations? Having analyzed the 10,000 randomly generated data sets, we found that for L1-L3, absolute stage durations were almost always more variable than relative stage durations, whereas for L4 the two tended to be the same (Figure 4C). The differences between variations of absolute vs. relative stage durations were more pronounced for empirical data, compared to permuted data. Adding all four stages of postembryonic development made this trend even more pronounced (Figure 4D).

**Figure 4.**
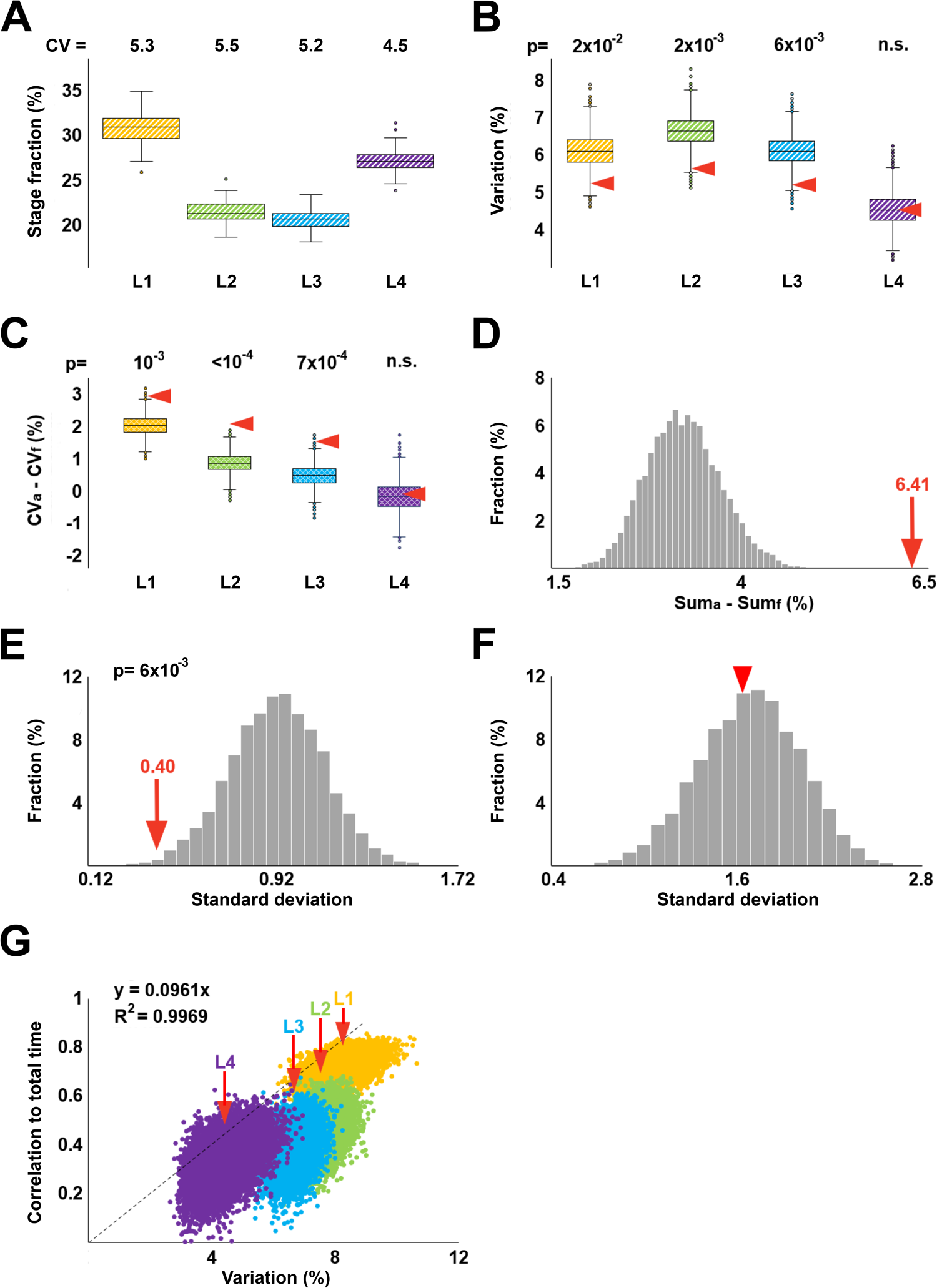
Fractional scaling of developmental time. **(A)** Fractional durations of larval stages calculated based on data in Figure 1D. Shown above are CVs (expressed as %) of L1-L4 stages. **(B)** Coefficients of variation of fractional stage durations from 10,000 randomly permuted sets. Arrowheads denote coefficients of variation of the empirical stage durations. **(C)** For each of the 10,000 randomly permuted sets, the CV of fractional durations (CVf) of a given stage was subtracted from the CV of absolute duration (CVa) of this stage; distributions of resulting values are represented as boxplots. Red arrowheads indicate CVa-CVf for the empirical data set. **(D)** The histogram in grey shows the following values for each of the 10,000 randomly permuted sets: (CVaL1 + CVaL2 + CVaL3 + CVaL4) – (CVfL1 + CVfL2 + CVfL3 + CVfL4). Red arrow indicates the same quantity obtained for the empirical data set. **(E)** Distribution of standard deviations of CVs of fractional stage durations from 10,000 randomly permuted data sets. p-value shows that only 58 of 10,000 values from permuted data are lower than the CV of fractional stage durations from the empirical dataset (red arrow). **(F)** Distribution of standard deviations of CVs of absolute stage durations from 10,000 randomly permuted data sets. Red arrowhead marks CV from the empirical dataset. **(G)** Pairs of *c*_*i*_, *s*_*i*_ values from each of 10,000 randomly permuted data sets compared to values of the empirical data set (indicated by red arrows). The four larval stages are depicted in different colors. The dashed line, the equation that describes it, and the R^2^ value are to demonstrate that *s*_*i*_ ≈ 10*c*_*i*_. In panels B, C, and E, p-values are the fractions of instances (out of 10,000) in which randomly permuted data were more extreme than those from the empirical dataset. In panel D, no value was as high as 6.41, indicating a conservatively estimated p < 10^−4^.

We also noted that unlike CVs of absolute stage durations that declined from L1 to L4 (Figure 1D), CVs of fractional times were quite similar. In fact, the standard deviation of these four values (5.28, 5.45, 5.17, 4.54%) was dramatically lower than the same quantity in the 10,000 data sets that were generated by randomly permuting stage durations of L1 through L4 (Figure 4E), whereas the same was not true for standard deviation of absolute stage durations (Figure 4F). Therefore, two observations are true – A) fractional times are less variable across individuals than absolute times and B) variability of fractional times is more similar across stages than variability of absolute times.

We examined the relationship between variation of absolute and fractional stage duration and found (Figure S4) that the two are related in the following way:

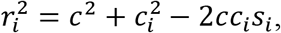

where *r*_*i*_ and *c*_*i*_ are CVs of fractional and absolute times, respectively, for *i*th larval stage; *c* is the CV of total (L1+L2+L3+L4) absolute time; and *s*_*i*_ is the correlation between absolute duration of *i*th larval stage and absolute total time. The above relationship is approximate and holds for *r*_*i*_, *c*, *c*_*i*_ ≪ 1.

It can be seen that if *c*_*i*_ = 2*cs*_*i*_, then *r*_*i*_ ≈ *c* < *c*_*i*_ and *r*_1_ ≈ *r*_2_ ≈ *r*_3_ ≈ *r*_4_. In the empirical data we analyzed, for all four larval stages, *c*_*i*_ = 2*cs*_*i*_ to within 16% of the value of *c*_*i*_, whereas in the 10,000 randomized data sets such modest deviations were essentially never observed (0, 0, 0, and 7 times for L1, L2, L3, and L4, respectively). The marked differences between the empirical and randomly permuted data are illustrated in Figure 4G – for all four larval stages, values of *c*_*i*_/*s*_*i*_ reside effectively outside respective permuted distributions. Moreover, the values of *c*_*i*_/*s*_*i*_ are similar for L1-L4 (*s*_*i*_ ≈ 10*c*_*i*_, see nearly linear relationship in Figure 4G), precisely as would be expected if *c*_*i*_ = 2*cs*_*i*_. This observation can account for variation in fractional time being lower than variation in absolute time and variation of fractional times being more consistent across different larval stages. It is not currently clear why *c*_*i*_ ≈ 2*cs*_*i*_. Our analysis suggests that *c*_*i*_ ≈ *cs*_*i*_ if durations of larval stages were perfectly correlated, whereas *c*_*i*_ ≈ 4*cs*_*i*_ if they were uncorrelated. The simplest interpretation of *c*_*i*_ ≈ 2*cs*_*i*_ is that stage durations are somewhat correlated. It will be interesting to determine whether the coefficient 2 is due to happenstance or a deeper, yet to be discovered relationships of stage durations.

## Discussion

Understanding mechanisms that regulate the temporal progression of development is an important problem, with much yet to be learned (Ebisuya and Briscoe, 2018). Species-typical average times are sufficient for addressing some questions, such as establishing timelines of specific developmental programs and studying molecular perturbations that alter them. Other mechanistic insights will require explicit consideration of inter-individual variation. Examples include testing whether developmental clock keeps absolute or relative time and understanding how timing of specific developmental events scales across individuals and environmental conditions.

We described a computational method that relies on minimal and apparently reasonable assumptions to infer duration of larval stages of individual *C. elegans* hermaphrodites from continuous measurements of their locomotor activity. One assumption that remains to be tested is whether our operational definition of stage boundaries (as midpoints of periods of reduced activity) yielded systematically different estimates of stage durations compared with ecdysis, which actually marks transition between stages. Our estimates of average stage durations are highly concordant with those obtained previously. Estimates of population-wide averages are robust and the sample of 125 individuals is ample for the task. However, although this is among the largest sets used for inferring developmental timing in *C. elegans*, future studies should be designed to be considerably larger to fully capture individual-to-individual variability and to infer stage correlations, both features being susceptible to the effects of outlying values. Precision of the method also requires that provisions must be made to explicitly account for batch effects that inevitably arise from multiple and difficult-to-control sources of variability. Our experience suggests that a reasonable tradeoff for larger sample sizes would be recording frequency ~0.1 Hz, which is more than an order of magnitude lower than those commonly used in studies of behavior.

Our analysis of individual timelines revealed several features of postembryonic development that could not have been identified if substantially fewer individuals were studied or if only population-average metrics were considered. These findings coalesce around three main ideas: First, the dramatic increase in body length (~4X; (Hirsh et al., 1976)) and volume (~10X; (Uppaluri and Brangwynne, 2015)), that occur during larval development in *C. elegans*, require voracious food consumption. Our analyses suggest that only during the L2 stage the rate of development is correlated with overall locomotor activity, which reflects foraging behavior (Calhoun et al., 2014; Flavell et al., 2013). This somewhat surprising result may imply that even in the animals that display the highest levels of exploratory activity, nutrient intake is sufficiently high to permit fast larval development. Alternatively, appropriate features of exploratory behavior that could predict developmental rate remain to be discovered as are environmental conditions that would make exploratory activity rate-limiting for postembryonic development. The final commitment to reproductive development (as opposed to dauer) occurs in L2 (Schaedel et al., 2012), which may require higher sensitivity to nutrition during this stage and thus help to explain the tighter coupling between activity and rate of development.

Second, there appears to be two separable phases during postembryonic development – one comprised of the first three larval stages (L1-L3) and the second of the L4. Absolute durations of L1-L3, unlike durations of the L4s, are significantly different between fast and slow developing animals (Figure 3C). Durations of L1-L3 are at least somewhat correlated to each other, but far less so to L4 (Figure 3D, E). Another way to illustrate the dichotomy between L1-L3 on the one hand and L4 on the other hand, can be seen in a comparison, using fractional times, of developmental progression of 20 fastest vs 20 slowest developers. Fractions of overall development time devoted to L1, L2, and L3 were indistinguishable between these two groups, whereas L4 distributions were essentially nonoverlapping (Figure 5A). This observation is opposite to what we found when analyzing the same data using absolute times (Figure 3C). The simplest hypothesis to account for these findings is that the dichotomy between L1-L3 and L4 reflects two different underlying processes, one for each of these two phases, that are at least somewhat decoupled. It is not currently clear what these processes might be, but an intriguing possibility is that duration of L1-L3 reflects some aspect of somatic development, whereas the L4 is dominated by germline production.

**Figure 5.**
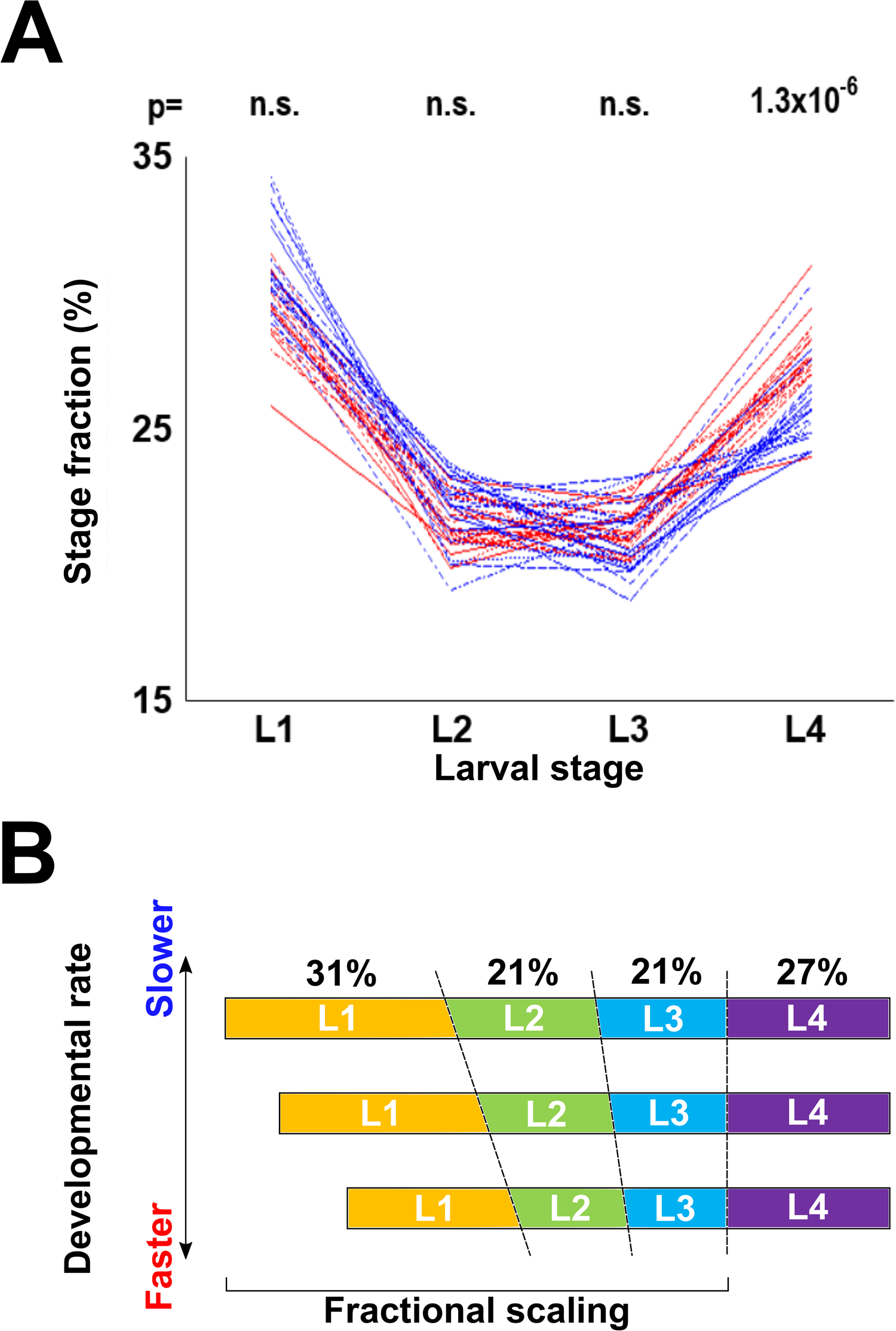
Temporal organization of postembryonic development in *C. elegans*. **(A)** Fractional stage durations of the 20 fastest and 20 slowest worms to reach adulthood. p-values shown above each stage are results of the Kolmogorov-Smirnov test comparing durations of that stage for the 20 fastest and 20 slowest developing worms. Compare with Figure 3C. **(B)** Model of temporal organization of postembryonic development in *C. elegans*. The three profiles shown correspond to a slow, intermediate, and fast developing individuals. Despite differences in absolute duration of development, fractions of overall developmental time devoted to each larval stage are conserved, primarily due to proportional scaling of L1-L3. Absolute duration of the L4 stage is less variable among individuals.

Third, some animals develop consistently faster than others at least in part because absolute durations of the first three larval stages are somewhat positively correlated (Figure 3D). We argue that these correlations arise from scaling of absolute durations of the first three larval stages with respect to the total duration of development. A plausible mechanism to account for this observation could be a “sizer” model that stipulates that in *C. elegans* molts occur once larvae reach a certain volume/weight (Uppaluri and Brangwynne, 2015), akin to size-related checkpoints that are critical in postembryonic development in insects (Nijhout, 2003; Rewitz et al., 2013). Developmental rates differ among animals, but a size constraint that is a fraction of a size required to attain adulthood would impose proportional scaling on developmental time, such that fractional durations would be relatively constant across individuals and environmental conditions. Absolute duration of the L4 stage shows little correlation with the three earlier stages, but it is more tightly constrained, therefore contributing less to the variability of the overall developmental time (Figure 5B). Our analyses suggest that a study of mechanisms that control scaling of L1-L3 stages, duration of the L4 stage, and relationship between variability and developmental timing, is likely to be fruitful.

## Materials and Methods

### Primary data and inference of stage durations

All primary data were generated and reported by Stern *et al.* (Stern et al., 2017). These data consisted of series of coordinates inferred from sequentially recorded frames that sampled, at 3 Hz, movement of individual animals. Each X-Y coordinate (~6 × 10^5^ per animal, spanning from L1 to adulthood) represents “center of mass” of an image of moving individual animal in a given frame (Stern et al., 2017). We obtained these coordinates from Mendeley (https://data.mendeley.com/datasets/3j6fsr634d/1). From coordinates corresponding to pairs of sequential frames, we calculated Euclidian distances that represented “displacements” over 1/3 second.

Because displacements between neighboring frames were a) highly variable and b) small with respect to animal size and average velocity, we experimented with effectively reducing recording frequency by calculating displacements between frames *n* and *n+x*, where *x* varied from 2 to 100 We refer to this process as reduction. We found that reduction to a sampling frequency of 0.1 Hz was a reasonable compromise between decreasing volatility of neighboring displacement estimates and retaining finer features of locomotor activity. Thus-obtained activity profiles were smoothened by averaging displacements in a sliding window of a fixed size that shifted by 1 displacement value at a time.

We operationally defined midpoints of periods of reduced locomotor activity as transitions between larval stages. Because shapes of activity profiles during periods of reduced activity were irregular and varied across stages (Figure 1C) as well as among worms, we developed an algorithm to estimate their width. We started by identifying four global activity minima per profile (*i.e.*, per worm), each corresponding to one period of reduced activity. Next, we drew 20 horizontal lines, first being 2.5 μm above the minimum and each subsequent one 2.5 μm above the previous line. Intersections of these lines with the activity profile defined the width of the period of lower activity at that vertical level. Finally, we averaged midpoints of these 20 width estimates thus obtaining provisional estimates of stage boundaries; these were further corrected in two ways.

First, the original data (Stern et al., 2017) contained missing frames incurred for technical reasons. Although there were few such frames (<0.4% of the total), we added their duration (1 frame = 1/3 second) to the estimates of stages during which they occurred. No extended runs of missing frames occurred sufficiently close to provisional boundaries between stages to meaningfully impact our ability to estimate them. Second, the algorithm that calculated smoothened activity profiles assigned value for each window based on the average of displacement values before (50%) and following (50%) it. Because this effectively shortened duration of the L1 by ½ of the sliding window size, we added this time to provisional estimates of duration of this stage.

### Computation of activity, correlations, and randomized developmental time series

We used two metrics to evaluate worm activity. First, we added all sequential displacements within a relevant stage. Activity defined in this way will be greater over longer time intervals. For this reason, there was a strong, but entirely uninformative correlation between activity and stage duration. We therefore computed correlations between stage duration and measures of activity that were normalized by stage duration. These latter quantities are equivalent to average velocities over the duration of a larval stage. Second, we calculated roaming fractions, as described previously (Ben Arous et al., 2009; Churgin et al., 2017; Flavell et al., 2013; Stern et al., 2017). Roaming fraction reflects the percentage (over a larval stage or entire postembryonic development) of behavioral episodes (each of a certain of defined duration) that were classified as roaming (as opposed to dwelling). For analysis in Figure 2C, activity profiles were manually classified into one of three categories (by two independent operators, with high concordance).

To evaluate several hypotheses against a null model of random association between stage durations, we generated 10,000 artificial data sets, each containing 125 developmental time series. Each time series was composed from an L1, an L2, an L3, and an L4, each randomly drawn from the set of empirically estimated values described in the section above.

### Statistical analyses

Data analyses were carried out using custom-written code (deposited to XYZ), Excel, and R package (https://cran.r-project.org/package=dgof). To evaluate variability across samples that in some instances had substantially different means, we routinely used coefficient of variation (CV), computed as standard deviation divided by the sample mean. Because standard deviation is susceptible to effects of outlier values, we used methods for comparing variation that were less affected by the extremes. We computed interquartile variability (based on Q2 and Q3 only), interdecile variability (2-9th deciles), or median absolute deviation of all data. In all cases, our conclusions regarding lower variability of L4 duration and less variable fractional times (compared to absolute times) held. One standard deviation of the correlation between two random sets of *N* values (expected to equal zero) is approximately 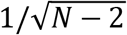, or ~0.09 for *N*=125. To account for multiple comparisons, we conservatively considered as significant only correlations >0.27 (three standard deviations), which correspond to p-values less than ~1.5×10^−3^. For comparing empirical data to results of permutation tests (using 10,000 randomly generated data sets) we considered as p-values the fractions of instances (out of 10,000) that were more extreme than the empirical values. In the box plots in all figures, the middle line is the median, top and bottom of the box encompass 2d and 3rd quartiles, and the whiskers represent the bulk of the fitted normal distribution.

## Acknowledgments

We thank R. Morimoto for generous hospitality and S. Stern for sharing data and advice. IR and VG are grateful to Marshall Butler. This work was funded in part by the NSF (IOS-1755244) and NIH (R01GM126125) grants to IR.

---

**Figure S1.**
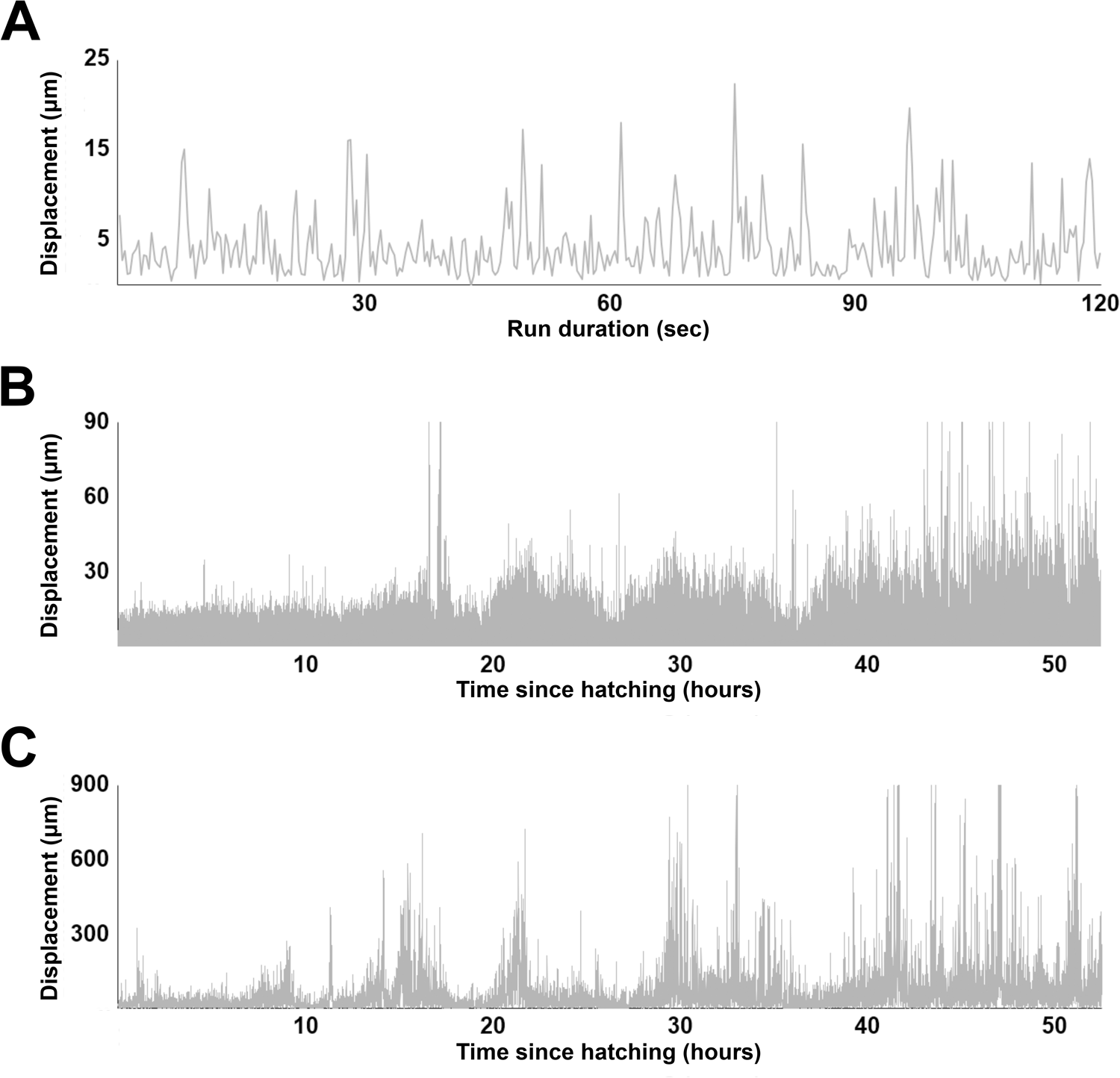
Variability of exploratory behavior. **(A)** A plot of 2 minutes (360 frames) of exploratory movement of the same animal as shown in Figure 1B. Note the average displacements of ~4-5 μm and that high displacement values (~15-20 μm) approximately correspond to the upper margin of much of the plot in Figure 1B. **(B)** Activity profile over the entire ~50 hours of recording of a different animal than the one shown in Figure 1B. Temporal pattern of displacements in this individual somewhat obscures periods of reduced activity corresponding to lethargus. **(C)** 30-fold reduced frame sampling still results in a noisy activity profile (this is the same animal as shown in Figure 1B), necessitating smoothing.

**Figure S2.**
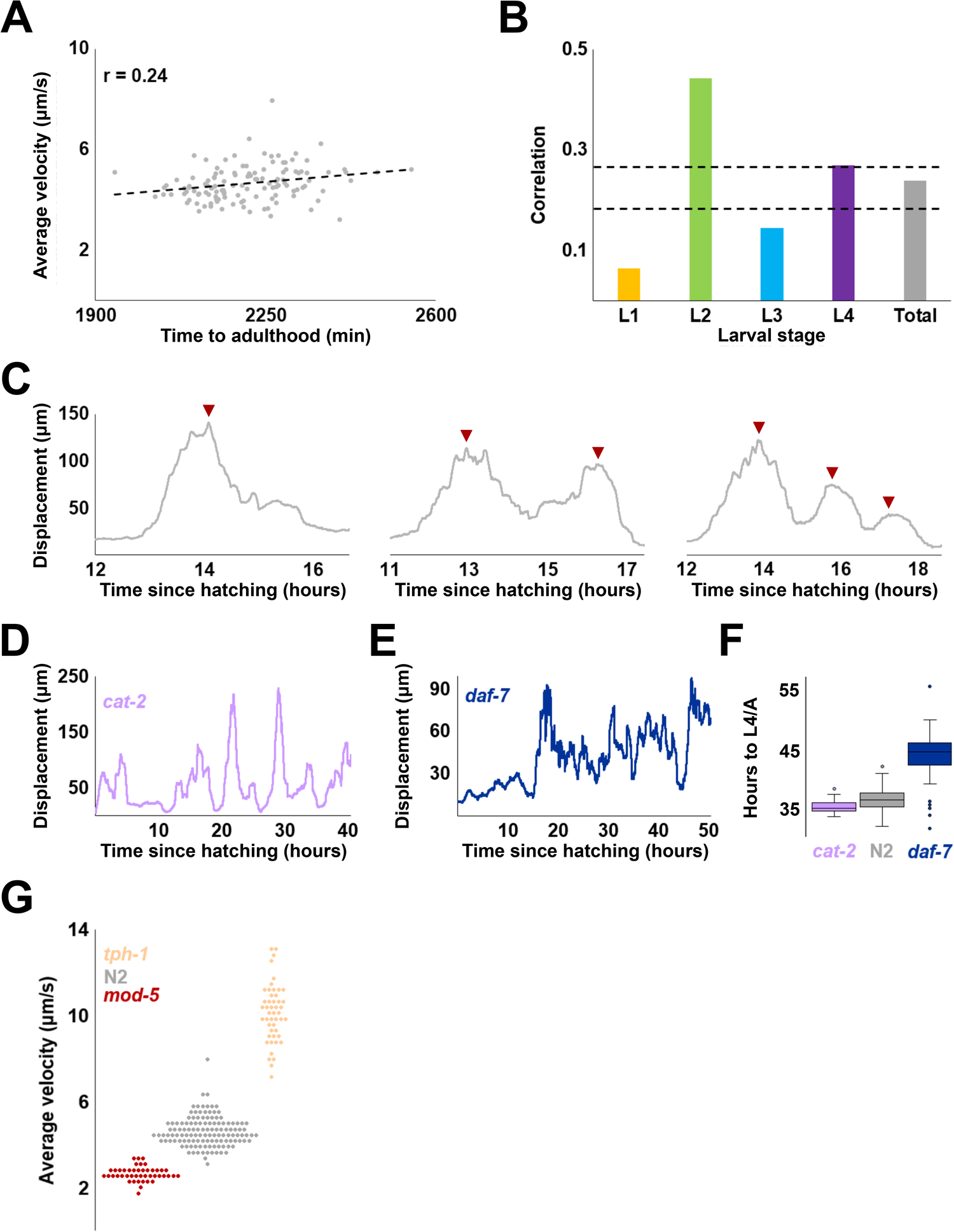
Relationship between locomotor activity and rate of development. **(A)** Correlation between total developmental time and locomotor activity (sum of displacements divided by time) for 125 wild type N2 worms. **(B)** Correlation between activity (sum of displacements divided by time) and stage duration for each larval stage; “total” shows the same value as in (A). Dashed lines denote two and three standard deviations above the expected correlation between two random sets of 125 uncorrelated variables. **(C)** Representative activity profiles of individuals representing 1 peak, 2 peak, and 3 peak (marked with arrowheads) categories. **(D)** Representative activity profile of *cat-2(e1112)* mutant individual. **(E)** Representative activity profile of *daf-7(e1372)* mutant individual. Note that inferring the L2/L3 boundary is particularly challenging. **(F)** Duration of development (*i.e.*, time to L4/adult transition) for *cat-2* (N=52) and *daf-7* (N=46) mutants compared to wild type N2 (N=125). **(G)** Total activity (sum of displacements divided by total time of development; this is equivalent to average velocity) for *tph-1*, N2, and *mod-5* animals. Each dot is one individual. Averaged activity profiles of these strains are shown in Figure 2D.

**Figure S3.**
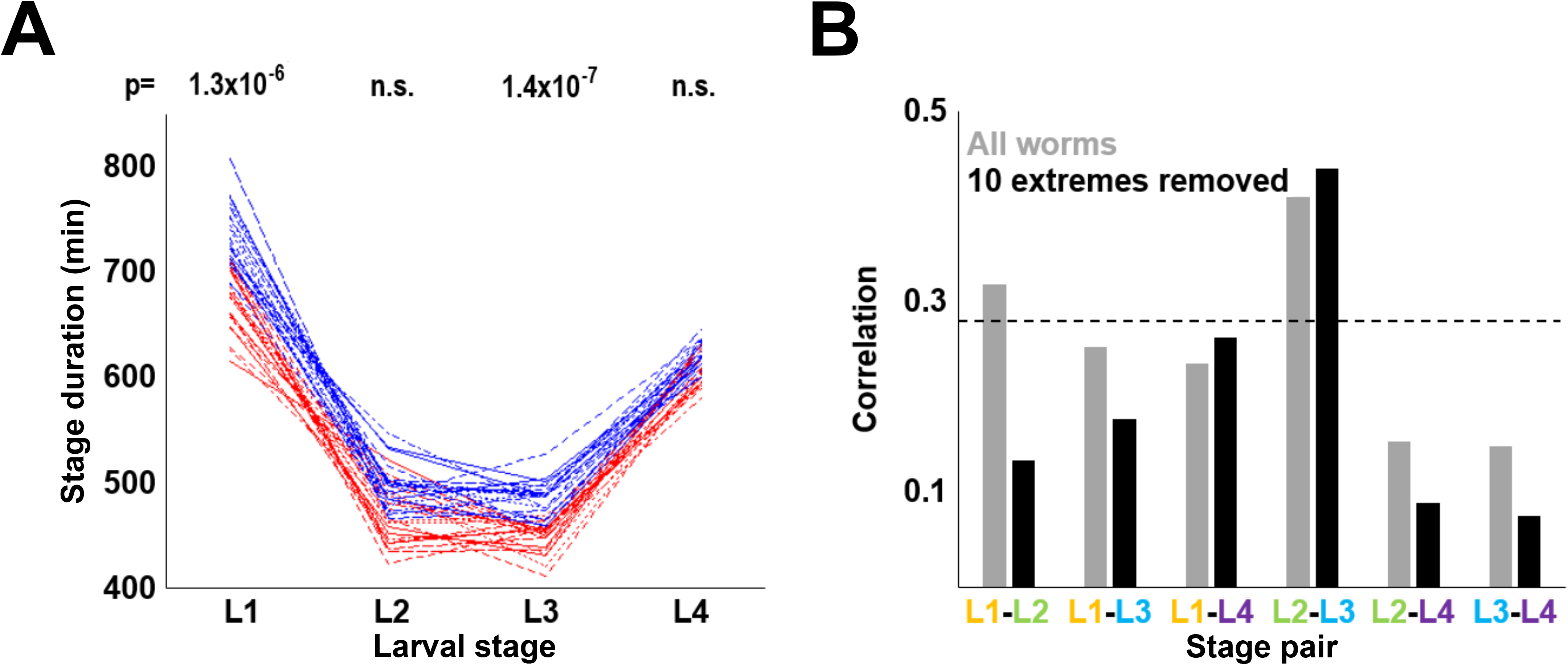
Sensitivity of analyses in Figure 3 to extreme values. **(A)** Stage durations of the 21st-41st fastest vs 21st-41st slowest worms to reach adulthood. p-values shown above each stage are results of the Kolmogorov-Smirnov test comparing durations of that stage for the 20 fastest and 20 slowest developing worms. **(B)** Pairwise correlations of stage durations of empirical N2 data including all 125 (in grey) or 115 (in black) individuals. To obtain the set of 115 from the set of 125, 10 individuals showing greatest deviations from population average developmental time were removed. Dashed line (0.27) denotes three standard deviations above zero, which is the expected correlation of two sets of random variables (N=125).

Durations of L1, L2, L3, L4 stages are *t*_1_, *t*_2_, *t*_3_, *t*_4_ respectively. *T* = *t*_1_ + *t*_2_ + *t*_3_ + *t*_4_. These quantities fluctuate about average values,

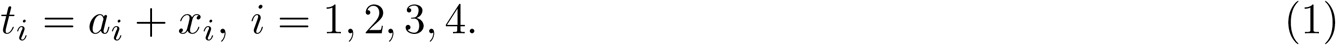

Here *a*_*i*_ > 0 are defined as

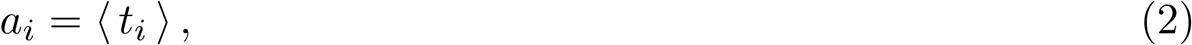

and the brackets ⟨ ⟩ stand for averaging. |*x*_*i*_| ≪ *a*_*i*_ are fluctuations. By definition, ⟨*x*_*i*_⟩ = 0.

Similarly, with the definitions *a* = *a*_1_ + *a*_2_ + *a*_3_ + *a*_4_ and *x* = *x*_1_ + *x*_2_ + *x*_3_ + *x*_4_, we find

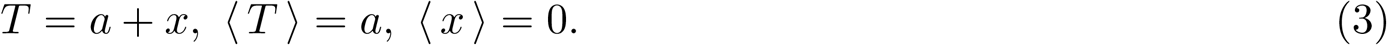

Define

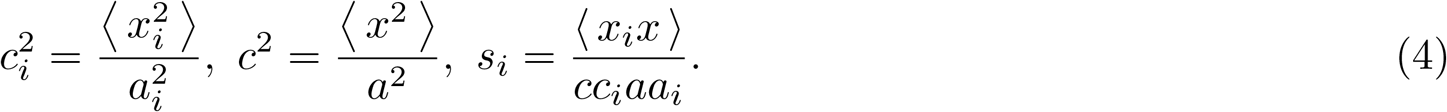

Here *s*_*i*_ are the correlation coefficients between the time of the *i*-th stage and the total time. Consider now the following quantities

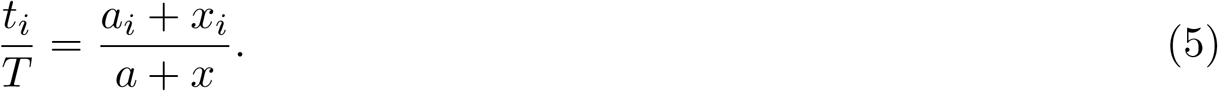

We would like to evaluate

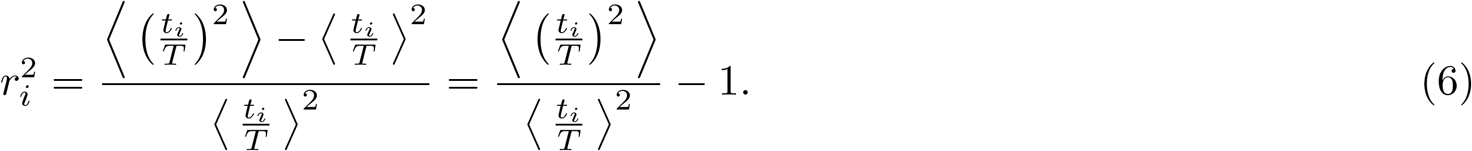

in the lowest order of perturbation theory in *x*_*i*_.

First we observe that

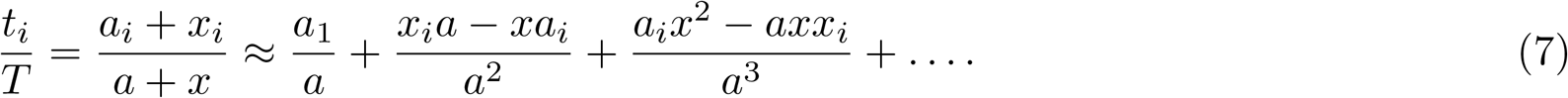

Averaging we find

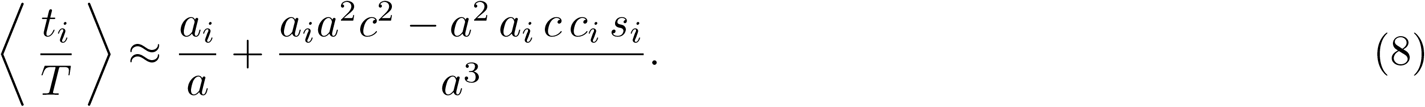

Now we expand

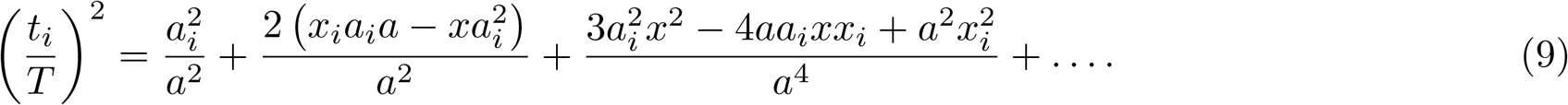

Averaging this, we find

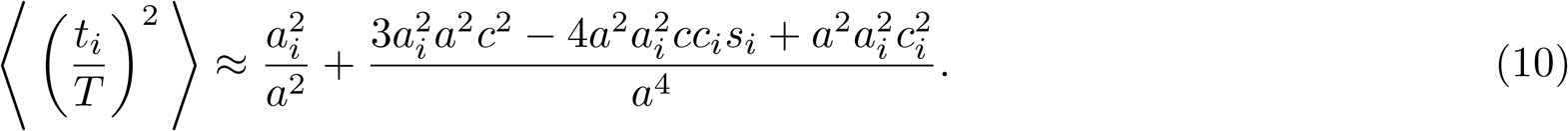

Finally, calculating the ratio and expanding, we find

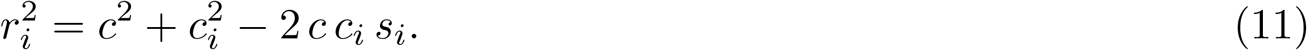

**Figure S4. Analytical derivation of the relationship between variation of absolute and fractional stage duration.**

